# Analysis of soybean germination, emergence, and prediction of a possible northward expansion of the crop under climate change

**DOI:** 10.1101/632976

**Authors:** Jay Ram Lamichhane, Julie Constantin, Céline Schoving, Pierre Maury, Philippe Debaeke, Jean-Noël Aubertot, Carolyne Dürr

## Abstract

Soybean (*Glycine max* (L.) Merr.) has potential to improve sustainability of agricultural production systems. A higher focus on this crop is needed to re-launch its production in the EU. A better understanding of key determinants affecting soybean establishment represents a first step to facilitate its adoption in cropping systems. To this objective, we conducted laboratory and field experiments in order to better characterize seed germination and seedling growth in relation to temperatures, water content, and soil structure. We then used these data to parametrize the SIMPLE crop emergence model and to evaluate its prediction quality, by comparing observed field germination and emergence data with the predicted ones. Finally, we performed a simulation study over the 2020-2100 period, for three sowing dates, from mid-March to mid-April, in the northern climate of France to evaluate whether future climate change will help expand soybean from Southern to Northern part of the country. Soybean germination was very fast, taking only 15 °C days to reach 50% germination at optimal conditions. The base, optimum and maximum temperatures were determined as 4, 30 and 40°C, respectively while the base water potential was −0.7 MPa, indicating a high sensitivity to water stress. The SIMPLE model well-predicted germination and emergence courses and their final rates, compared with the observed field data. The simulation study showed average emergence rate ranging from 61 to 78% with little variability among sowing dates and periods, but a high variability between years. Main causes of non-emergence were seedling mortality due to clods or soil surface crust followed by non-germination and seedling mortality due to drought, especially for mid-April sowing. These results provide a better knowledge of soybean establishment that are encouraging to introduce soybean with early sowings to diversify current cropping systems.

## 1. Introduction

Crop diversification is one of the three measures of the EU common agricultural policy greening initiative that aims to enhance the environmental performance of the EU agricultural system (European Commission, 2013). Leguminous crops in arable rotations are encouraged to this objective as they provide multiple ecosystem services (Schreuder and De Visser, 2014; Stagnari et al., 2017).

Soybean (*Glycine max* (L.) Merr.) is one of the leguminous crops with a strong potential to improve sustainability of agricultural production systems. Despite this potential, a great decrease of the soybean acreage has been observed in the EU since 2002 which was mainly due to an insufficient economic competitiveness of the crop compared with non-leguminous crops (FAOSTAT, 2018; Labalette et al., 2010). A reintroduction of soybean is one of the public policy priorities both to reduce import dependency of this crop and also to satisfy the demand for locally produced, non-genetically modified protein crops (Bertheau and Davison, 2011). In France, soybean surface grown has been increasing over the last years and the crop is mainly grown in east-central and southwestern regions. Nevertheless, soybean is already grown to the north of the Paris basin, where the surface area has increased from 30 to 340 hectares in the last four years (Lejeune-Henaut, 1991; Ribault, 2018). There is an effort to increase the competitiveness of this crop at the expense of cereals by maximizing quantitative and qualitative yield per unit area. A better understanding of key biotic and abiotic factors that affect crop establishment is crucial and represents a first step to this objective.

Soybean is a spring crop mainly grown across French regions characterized by low annual rainfall. While no important losses due to biotic stresses have been reported to date, cultivation across southern regions often poses risks related to the seedbed abiotic components that affects the crop establishment. Indeed, stand establishment of field crops is widely affected by soil temperature, water potential, as well as mechanical obstacles such as soil aggregates composing the seedbed or a soil surface crust (Awadhwal and Thierstein, 1985; Constantin et al., 2015; Dürr et al., 2016; Dürr and Aubertot, 2000; Gallardo-Carrera et al., 2007). Nevertheless, only little knowledge exists to date concerning the impact of seedbed abiotic components on soybean crop establishment.

Three were the objectives of this study. First, get a better knowledge of soybean parameter values with regard to germination and early seedling growth in different environmental conditions. Second, use these data to parametrize the SIMPLE crop emergence model for soybean, and evaluate the prediction quality of the model -- in terms of germination and emergence courses and their final rates -- by comparing the predicted and observed data under field conditions. Third, use this model adapted to soybean to perform a simulation study that helps analyze whether there will be suitable seedbed sowing conditions for soybean in Northern France under future climate change. This simulation study is important given that a global northward expansion of agricultural climate zones has been predicted under 21^st^-century climate change (King et al., 2018). A total of 243-soybean emergence simulations was performed taking into account three sowing dates, from mid-March to mid-April. To this aim, we first mobilized the STICS soil-crop model (Brisson et al., 2003, 1998) to generate soil water content and temperature in the seedbed (0–10 cm) using the most pessimistic IPCC scenario. We then used the data obtained as input variables to feed the SIMPLE crop emergence model (Constantin et al., 2015; Dürr et al., 2001). The emergence courses and final rates, and causes of no-seedling emergence are analyzed and discussed.

## 2. Materials and methods

### 2.1. Overview of the SIMPLE crop emergence model

A comprehensive description including the functioning of the SIMPLE model and the list of equations and input variables has been previously provided (Dürr et al., 2001). Briefly, the model predicts the germination and emergence process and their final rates in relation to environmental conditions during sowing. The model has previously been parameterized and positively evaluated for a number of crop species: wheat, sugar beet, flax, mustard, French bean, oilseed rape, (Dorsainvil et al., 2005; Dürr et al., 2016, 2001; Moreau-Valancogne et al., 2008), several catch crops (Constantin et al., 2015) and a plant model *Medicago truncatula* (Brunel et al., 2009). Here we mainly focus on the model’s key features and the input variables measured for the model parametrization for soybean crop without listing equations.

SIMPLE creates 3D representations of seedbeds with sowing depth distribution and the size, number, and position of the soil aggregates as input variables. Soil temperature at the mean sowing depth and daily soil water potential in several layers are also used as input variables for simulations, along with plant characteristics for germination and seedling growth. Seeds are placed into the seedbed at random using the sowing depth distribution chosen for the simulation. The model predicts germination and emergence, seed by seed, at a daily time step. The time required for germination of the seed i is chosen at random in the distribution of thermal times that characterizes the seed lot used. Cumulative thermal time from sowing is calculated above the base temperature (Tb) for germination, provided that the soil water content at the seed sowing depth is above the base water potential (Ψb). The Tb and Ψb for germination are input variables. If seed i has not germinated after a given time (often fixed at 30 days for the simulation), the model considers it will never germinate. If the seed germinates, then a seedling grows from the seed. To better include the effect of early water stress on seedling growth, we added a water stress function to the SIMPLE model, which reduces emergence after germination (Constantin et al., 2015). With this function, the fate of seedlings is determined by considering soil water potential in the soil layer in which the radicle grows in the two days following germination. If it is lower than Ψb, the seedling does not emerge and dies the following day. If this is not the case, the time it takes for the seedling to reach the soil surface after germination is calculated by SIMPLE based on its seed’s sowing depth, the length of the pathway the shoot takes through the aggregates, and the shoot’s elongation function, whose parameters are input variables. The probability of the seedlings remaining blocked under aggregates depends on the size and position of the clods in the seedbed, i.e. lying on the surface or below it. Soil surface crusting depends on cumulative rainfall after sowing; a proportion of seedlings remain blocked under the crust depending on daily crust water content (dry or wet soil surface). Simulations are run for 1000 seeds to predict the emergence rate over time and final emergence percentage. The causes of non-emergence simulated by SIMPLE are (i) non-germination, (ii) death of seedlings caused by water stress after germination and (iii) mechanical obstacles (clods or a soil crust). The SIMPLE model does not consider biotic stresses, such as pests and diseases or the effect of high temperatures, which could inhibit germination.

### 2.2. Input variables of the model as determined by laboratory experiment and measurements

Laboratory experiments were carried out to establish reference values on soybean germination, especially in water stress conditions, which are lacking in the literature, and to get the input values necessary to parametrize the SIMPLE model.

#### 2.1.1. Seed germination

Seed germination tests were carried as previously described (Gardarin et al., 2016; Raveneau et al., 2011). A commercial seed lot of cv. ES Pallador, which was produced under good standard growing conditions in France, was used. No seed treatment was applied prior to sowing. For each temperature, four replicates of 25 seeds were sown in 90 mm Petri dishes on Whatman® filter paper of the same size placed both below and above the seeds and imbibed with 8 ml deionized water. The dishes were incubated at 3.5, 6.5, 10, 15, 20, 25, 30, 34.5, 37 and 40 °C and temperatures were recorded hourly with sensors. Depending on the incubated temperature, seed germination was assessed up to three times per day until no further germination was observed. A given seed was considered germinated when the radicle was >3 mm. At each observation, the germinated seeds were removed from the dishes. The base temperature for germination was determined by fitting a Gompertz function to the observed germination rates as described previously (Brunel et al., 2009). Adjustments of germination dynamics were made for each temperature and values of Tb for germination were defined as the X-intercept of the linear regression between the temperature and germination rate (Dahal and Bradford, 1994; Gummerson, 1986). We determined the range of temperatures for which a strong linear relationship existed between germination rates (1/time to reach 20, 40, 50, 60, 80, and 90% germination) to calculate the X-intercepts. Base temperature was defined as the intercept with the X-axis, temperature fitted value at which no germination occurs. The optimum temperature for germination, corresponding to the maximum germination speed, was determined using a non-linear equation (Yin et al., 1995), where maximum and minimum temperatures were found as function parameters.

Germination was also tested under four different water potentials (0, −0.10, −0.25, 0.50, − 0.75, −1.0 and −1.50 MPa) by adapting the previously described methods (Brunel et al., 2009; Gardarin et al., 2016; Raveneau et al., 2011). The same number of seeds per replicate as described above was used. Because the use of polyethylene glycol (PEG) solution delay seed germination (and increase the length of incubation), the seeds were disinfected by using 1% Sodium hypochlorite solution for 10 min followed by two rinses in distilled water. In preliminary tests, non-disinfected seeds were contaminated by mold beginning from 5 days post-incubation at 20°C that hindered the follow-up of measurements. Twenty-five disinfected seeds were laid onto flat Whatman filter paper in 90 mm Petri dishes with 20 mL of osmotic solutions of high molecular weight PEG (Polyethylene glycol 8000, ref. SIGMA 25322-68-3). The latter was used at different concentrations to control water potential as described previously (Michel, 1983). The quantity of PEG solution used was higher (20 ml) than the quantity of deionized water used (8 ml) since time to reach the maximum germination is longer with PEG solution. The same adjustments as described above were used to calculate base water potential for germination.

#### 2.2.2. Hypocotyl and radicle elongation

Destructive measurements of hypocotyl and radicle growth under dark (heterotrophic growth) were performed adapting previously described methods (Brunel et al., 2009; Durr and Boiffin, 1995). Seeds were sown at a depth of 3 cm in polyethylene pots (five seeds per pot; 30 pots in total; ⌀ = 10.5 cm; h = 11.5 cm) filled with 500 ml white sand (150-210 μm). The pots were leaned on underlying plastic pots. The moisture gravimetric content was raised to 0.20 g g^−1^ before sowing by watering the sand with 100 ml of distilled water. The pots were covered with aluminum foil, to avoid light penetration and prevent water loss, and incubated in a dark growth chamber at 20 ° C and 80% relative humidity.

Seeds and seedlings were harvested by sampling three pots every day beginning from 3 days after sowing (das). At each observation (10 in total), 15 seedlings were recovered. Plant organs were separated from the sand, washed and radicle and hypocotyl lengths were measured. Results were fitted to a Weibull function for hypocotyl and a probability function for radicle.

#### 2.2.3. Seedling death under soil aggregates

This test was performed in a growth chamber as described previously (Dürr and Aubertot, 2000). Soil was sampled from 0-20 cm soil horizon of a field plot (the same where field experiment was conducted), sieved through 5 mm holes and stocked in sealed plastic containers until its use to avoid water loss. The gravimetric water content of the soil sample was measured as described previously (Gallardo-Carrera et al., 2007). Aggregates of different sizes were collected from the same field plot and air-dried for four weeks. The aggregates were grouped in three classes according to their longest axis L: 30-40, 40-50 and 50-60 mm.

Two experiments were performed: first with aggregates completely buried under soil and second with aggregates on the soil surface. In each experiment, three replicates per class of aggregate (24 aggregates/replicate = 72 aggregates/class, each replicate represented by a plastic tank, were used. Four liters of the field soil were placed on the bottom of each plastic tank (42 x 30 x 10 cm^3^) to create a first layer (Ca. 4 cm depth). Twenty-four seeds were sown on this soil layer using a grid to define their position in the tank. Another two liters of the soil was placed over the seeds to make a second layer (Ca. 2 cm depth) of the soil. For each class, aggregates were positioned on top using another grid, so that their centers were just above the seeds. The aggregates were then covered with a third (last) layer of soil (Ca. 3 cm), until their complete burial. This final layer of soil was not added in experiments with aggregates laid on the soil surface. All plastic tanks were carefully sealed with aluminum foil, to avoid light penetration and water loss, and were incubated in the dark at 20°C and 100% relative humidity for two weeks until when the maximum length of the hypocotyl was reached. At two weeks post incubation, each seed or seedling was observed in a vertical plan and its state was recorded: germinated or not, abnormal without striking an obstacle, blocked under an aggregate or not. The percentage of seedling blocked under aggregates (buried and on the soil surface) observed was fitted to a probability function as described previously (Dürr and Aubertot, 2000).

### 2.3. Field experiments

#### 2.3.1. Experimental site and climatic data

The experiment was carried out in May 2018 in Auzeville experimental station of INRA (43.53°N, 1.58°E). The soil had the following soil granulometry and chemical characteristics at the 0-30 cm soil horizon: 0.253 g.g^−1^ clay, 0.380 g.g^−1^ silt and 0.366 g.g^−1^ sand, 0.0914 g.g^−1^ C, 0.001 g.g^−1^N, C/N ratio 8.31, and pH 8.57, organic matter 0.009 g.g^−1^, bulk density at sowing depth 1.0 g.cm^−3^. After the harvest of wheat in July, conventional tillage was performed with 4-body plough in October 2017 at 30 cm depth. A 9-cm tillage was followed with seedbed cultivator (Kongskilde, tractor New Holland 115) on 13 March 2018 to prepare the seedbed. Soybean (cv. ES Pallador) was sown in 72 m^2^ blocks (each of 12 m long and 6 m wide, 4 blocks in total) on 22 May 2018 at 3 cm sowing depth with 40 seeds m^−2^ and with 50 cm inter-row distance.

Soil temperature and water content were recorded using climate sensors (ECH O 5TM, METER Group, Inc. USA). The sensors were installed in the seedbed following sowing at three soil horizons (3 sensors/depth/block at −3 cm, −5 cm and −10 cm). These sensors delivered hourly temperatures, measured by an onboard thermistor, along with accurate volumetric water content. Data were recorded from sowing until crop emergence. Rainfall data were obtained from an automatic meteorological station installed at the experimental site. The cumulative degree-days (°Cd) were calculated from sowing (time 0) as the sum of the average daily air temperature minus base temperature of soybean.

#### 2.3.2. Seedbed characterization

The distribution of seedbed aggregate size was characterized as described previously (Aubertot et al., 1999; Gallardo-Carrera et al., 2007). Seedbed samples were taken just after sowing. A surface was delimited along the row of the seedbed with combs (20 x 10 x 10 cm^3^) to determine the numbers of aggregates in a precise soil volume. The sample surface was painted with an aerosol bomb, to ensure complete cover to distinguish aggregates visible on soil surface (painted) from those buried within the seedbed (not painted). Four samples per replicate block (16 samples in total) were carefully extracted with a spoon from four different rows, brought to the laboratory, and dried in an oven at 105 °C for 24 hours. The samples were sieved with a gently shaking machine (30 s, 50-mm amplitude) and grades <5, 5-10, 10-20, 20-30, 30-40, 40-50, and >50 mm were obtained. Painted and non-painted aggregates were separated, weighed and counted. Three classes of aggregate burial were assessed visually from the painted portion of the aggregate surface: completely buried aggregates (no painted surface), partially buried (≤ 50 % of painted surface), and no buried aggregates (≥ 50% of painted surface).

Since soil water content – transformed into water potential – was the input variable of the SIMPLE model, sieved soil (5mm) from the same field plot was introduced into Richards’press to determine soil water-holding capacity at several pressures : −0.1, −0.3, −1.0, −1.3 and −1.5 MPa. Three replicates for each pressure were made and the relationship between soil water content and water potential was fitted to van Genuchten equation (van Genuchten, 1980).

#### 2.3.3. Measurement of sowing depth

The sowing depth was determined by performing a semi-destructive measurement the day after sowing. A micro-profile was made by moving vertically until reaching seeds along the row. Once seeds were retrieved, the sowing depth (i.e. the distance between the seed and soil surface) was measured by using a ruler. The measurement was performed for 100 random seeds/block (four replicates of 25 seeds along four rows in diagonal) for a total of four blocks (i.e. 400 seeds).

#### 2.3.4. Seed germination and seedling emergence rates

The number of germinated seeds was counted every day beginning from 2 das. The same method, as for sowing depth, was used on the same number of seeds. Seeds were considered to have germinated successfully if the radicle was > 0.5 cm in length. The measurements were continued until reaching a plateau (the same number of germinated seeds for three consecutive days) and when no rainfall was foreseen in the coming days. However, the measurements continued when rainfall was foreseen as it triggers germination of those seeds which did not germinate due to water stress.

Seedling emergence was counted in 4 m^2^/block (2 linear meters/ row, 4 rows in diagonal, for a total of 16 m^2^/plot) delimited with plastic pegs. A seedling was considered emerged when cotyledons were clearly visible over the soil surface (Fehr and Caviness, 1977). The counting was continued until reaching a plateau with no new emerging seedlings during several days.

#### 2.3.5. Causes of non-emergence

The causes of non-emergence were identified using a visual diagnostic key. Each symptoms or characteristics observed in the field was assigned to a specific stress as follows: i) no seed found: technical problem of sowing or predation of seeds by birds, ii) empty seed coat: damage caused by granivores, iii) intact seed without necrosis/rotting (N/R): pre-germination abiotic (water or temperature) stress, iv) seed or seedling showing N/R: pre-emergence damping-off, v) presence of holes or larvae in or around seeds/seedling: soil pests (seed maggot, wireworm etc.), vi) seeds germinated, no N/R, no aggregate above, but presence of soil crust: stress due to soil crust, vii) seeds germinated, no N/R, no soil crust but presence of aggregates above: seedling blocked under aggregates, and viii) seeds germinated, drying seedling, no N/R, no soil crust or aggregates, no or little rainfall after sowing: post-germination water stress.

To this objective, 60 empty points without seedling emergence/block (15 along the row, 4 rows in diagonal) were randomly selected. A micro-profile was made by a spoon at each empty point moving vertically until reaching non-germinated seeds or non-emerged seedlings. Once seeds or seedling parts were found, their status was annotated according to their characteristics and possible causes of non-emergence were noted.

### 2.4. Statistical criteria used for model evaluation

Three statistical criteria – model efficiency (EF), root mean square error (RMSE) and coefficient of residual mass (CRM) – were calculated to assess the quality of model predictions for germination and emergence.

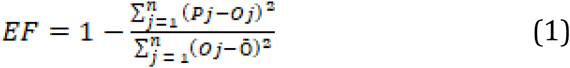

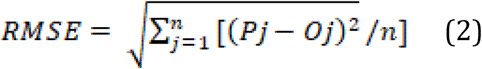

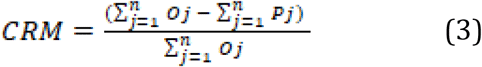

Where EF (ranges from −∞ to 1) represents model accuracy relative to the mean of observed data and is = 1 for a perfect model prediction. The more EF approaches 1, the more is the match between observed and predicted values. Pj and Oj are predicted and observed values, respectively, n is the number of observations, and Ō is the mean of observed values.

RMSE provides absolute error (the mean difference between n predicted and observed values). The unit of the coefficient is the same as that of the analyzed variables.

The coefficient of residual mass (CRM) is a measure of the tendency of the model to under-or overestimate predicted values compared with observations. A negative value indicates that the majority of predicted values are less than the observed ones (i.e. predicted time courses are ahead of the observed ones).

ANOVA was applied to determine any differences in mean values between treatments and their comparisons. All statistical analyses were applied using software R (Hothorn and Everitt, 2009).

### 2.5. Simulation study

#### 2.5.1. Climate scenarios and simulations of the seedbed climate

We used the RCP 8.5 emission scenario based on the regionalized climate projections data provided by Drias platform (www.drias-climat.fr). These data were used to generate soil temperature and water content of the seedbed using the STICS soil-crop model (Brisson et al. 2003). This model daily simulates soil water contents and temperatures, according to daily weather and soil characteristics. Variations in soil moisture were predicted using STICS at 0-2, 2-4, 4-6 and 6-10 cm. We chose Estrées-Mons (49°52′44″N 3°00′27″E) located in Northern France as representative area of North European climate. A detailed description of the study site and soil granulometry and chemical characteristics have been previously published (Lamichhane et al., 2019). The predicted daily values were used to feed the SIMPLE model.

#### 2.5.2. Soybean sowing scenarios

A coarse seedbed, the same as the one from our field experiment, and typical of that prepared by growers, was considered. This seedbed had 33 % of the soil aggregates >20 mm in diameter and 77% of its aggregates <20mm in diameter. The simulated sowing depths were 3 ± 1.5 cm.

A total of 243-soybean emergence simulations were performed for a period between 2020 and 2100, taking into account three sowing dates: mid-March, 1^st^ April, and mid-April. Although farmers in Northern France currently practice late April sowing, we included earlier sowings taking into account a possible shift in future sowing dates due to climate change.

#### 2.5.3. Analysis of simulation results

Data were pooled and analyzed by sowing date for each 20-year period. Twenty years considered for each period were treated as replicates. ANOVA was used to determine the potential effect of sowing dates for each period and their interaction both on soil weather and crop germination and emergence parameters.

The variability of germination and emergence rates and their duration were analyzed as described previously (Lamichhane et al. 2019). To this aim, we established three classes of rate or duration, expressed as the frequency of each class over the 20-year period for germination and emergence rates, and their duration. For germination rate, thresholds were poor germination when germination rate was <75% and sufficient germination above 75%. For emergence rate, thresholds were poor emergence when the emergence rate was <50%, and sufficient over 50%. For germination duration, thresholds were low duration when the number of days required to reach maximum germination (NGmax) was < 14 days and high when NGmax was >14 days. For emergence duration, thresholds were low duration when the number of days required to reach maximum emergence (NEmax) was < 28 days, and high when NEmax was >28 days. The frequency of poor germination (<75%) and emergence (<50%) rates as well as high NGmax (>14 days) and NEmax (>28 days) duration were analyzed as they could lead to crop emergence failure and potential re-sowing.

The variability of causes of non-emergence was analyzed by establishing two classes of seed and seedling mortality rates for each mortality cause. For non-germination, the two classes were low with <25% and high with >25% non-germinating seeds. For seedling mortality due to clod, crust and drought, the two classes were low with <15%, and high with >15% of seedling mortality. Frequency of high risks of non-germination (>25%) and seedling mortality due to clod, crust, and drought (each >15%) cases are presented for the same reason as described above. To determine significant effects on germination, emergence rates, and duration as well as on causes of non-emergence, we used the same statistical analysis performed for weather data. All statistical analyses were applied using software R (Hothorn and Everitt, 2009).

## 3. Results

### 3.1 Laboratory experiments

#### 3.1.1. Seed germination under laboratory conditions

The optimum temperature for soybean was 30°C and germination speed strongly decreased over this temperature **(Fig. 1a)**. The calculated base temperature for seed germination was 4 °C. The germination curve was then expressed as a function of thermal time (**Fig. 1b**) which finely grouped the germination results obtained in the linear part of Figure 1a (i. e. 10, 15, 20, 25 °C). A Gompertz function was fitted to these results and classes of germination times were created as input values for the SIMPLE model **(Table 1)**. Over 98% seeds germinated very rapidly in only 35 °Cd and time to reach 50% germination corresponded to 15 °Cd (**Fig. 1b**). When grouped by classes of °Cd, 8% of seeds germinated within the class 5-10 °Cd, followed by 51, 29, 8 and 2% of seeds germinated within the class 10-15, 15-20, 20-25, and 25-35 °Cd, respectively.

**Table 1.**
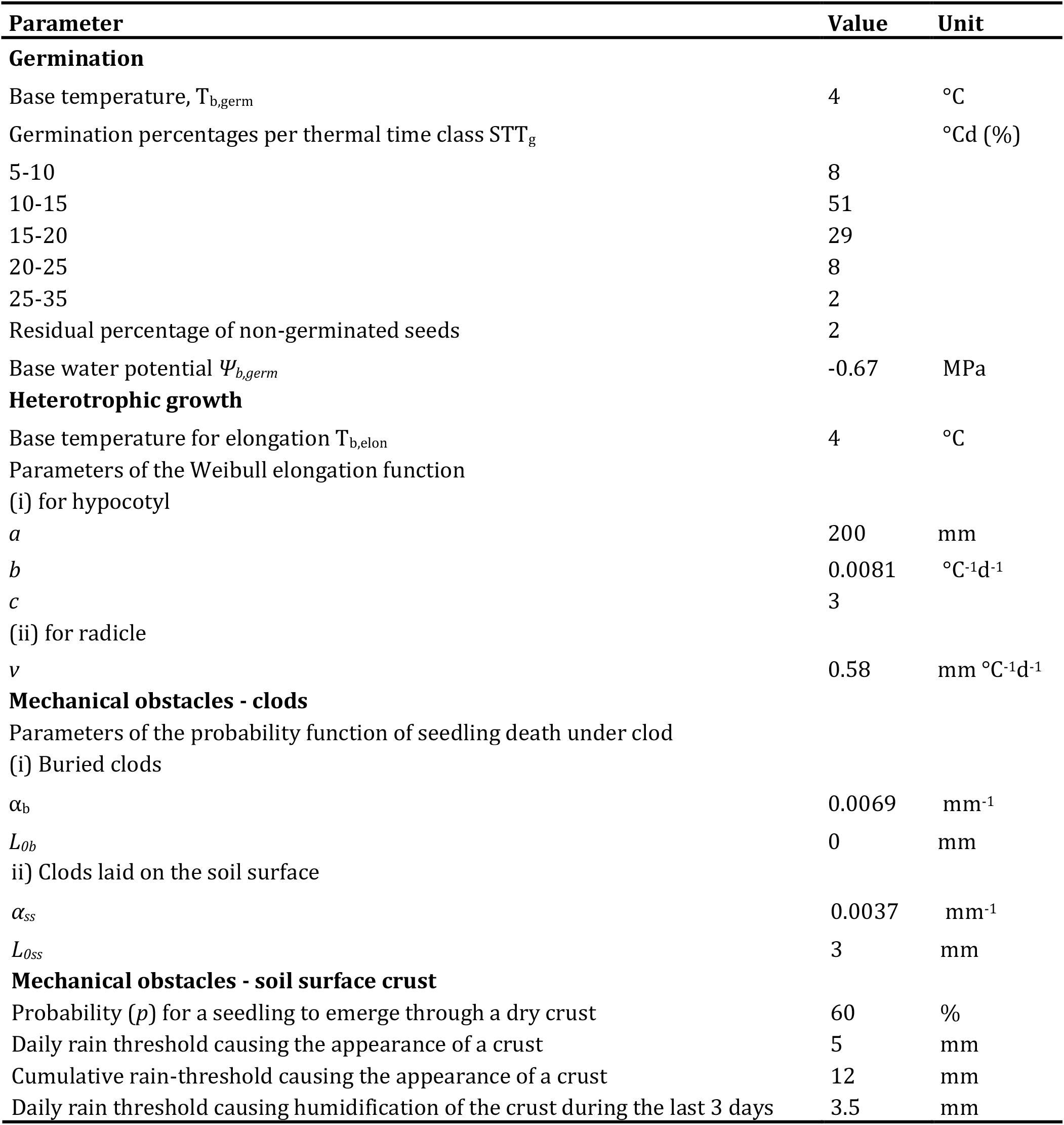
Values of the input variables of SIMPLE for soybean used in this study

| Parameter | Value | Unit |
| --- | --- | --- |
| **Germination** |  |  |
| Base temperature, $T_{b,germ}$ | 4 | °C |
| Germination percentages per thermal time class $STT_g$ | | °Cd (%) |
| 5-10 | 8 |  |
| 10-15 | 51 |  |
| 15-20 | 29 |  |
| 20-25 | 8 |  |
| 25-35 | 2 |  |
| Residual percentage of non-germinated seeds | 2 |  |
| Base water potential $\Psi_{b,germ}$ | -0.67 | MPa |
| **Heterotrophic growth** |  |  |
| Base temperature for elongation $T_{b,elon}$ | 4 | °C |
| Parameters of the Weibull elongation function |  |  |
| (i) for hypocotyl |  |  |
| $a$ | 200 | mm |
| $b$ | 0.0081 | °C<sup>-1</sup>d<sup>-1</sup> |
| $c$ | 3 | |
| (ii) for radicle |  |  |
| $v$ | 0.58 | mm °C<sup>-1</sup>d<sup>-1</sup> |
| **Mechanical obstacles - clods** |  |  |
| Parameters of the probability function of seedling death under clod |  |  |
| (i) Buried clods |  |  |
| $\alpha_b$ | 0.0069 | mm<sup>-1</sup> |
| $L_{ob}$ | 0 | mm |
| ii) Clods laid on the soil surface |  |  |
| $\alpha_{ss}$ | 0.0037 | mm<sup>-1</sup> |
| $L_{oss}$ | 3 | mm |
| **Mechanical obstacles - soil surface crust** |  |  |
| Probability ($p$) for a seedling to emerge through a dry crust | 60 | % |
| Daily rain threshold causing the appearance of a crust | 5 | mm |
| Cumulative rain-threshold causing the appearance of a crust | 12 | mm |
| Daily rain threshold causing humidification of the crust during the last 3 days | 3.5 | mm |

**Figure 1.**
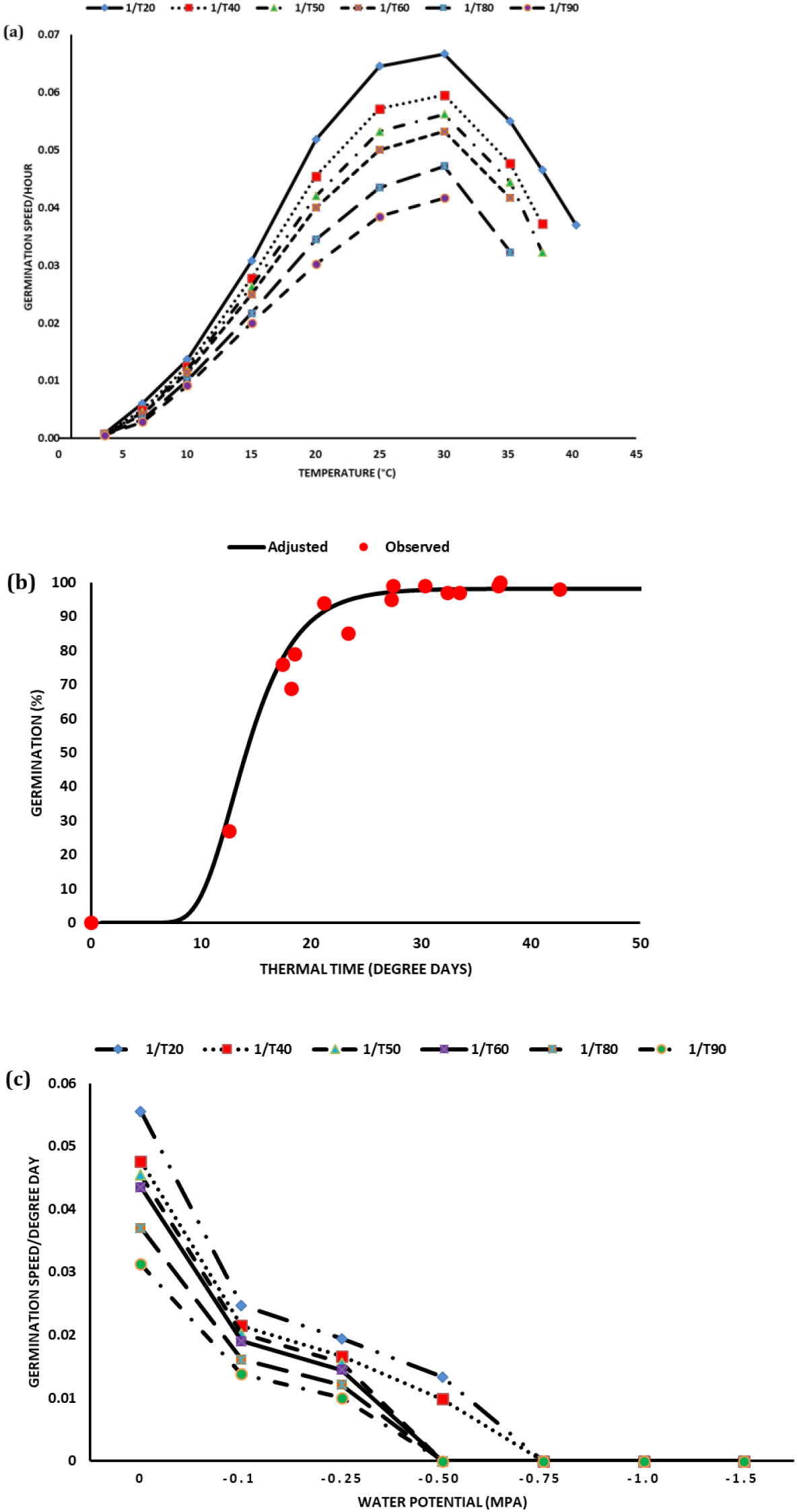
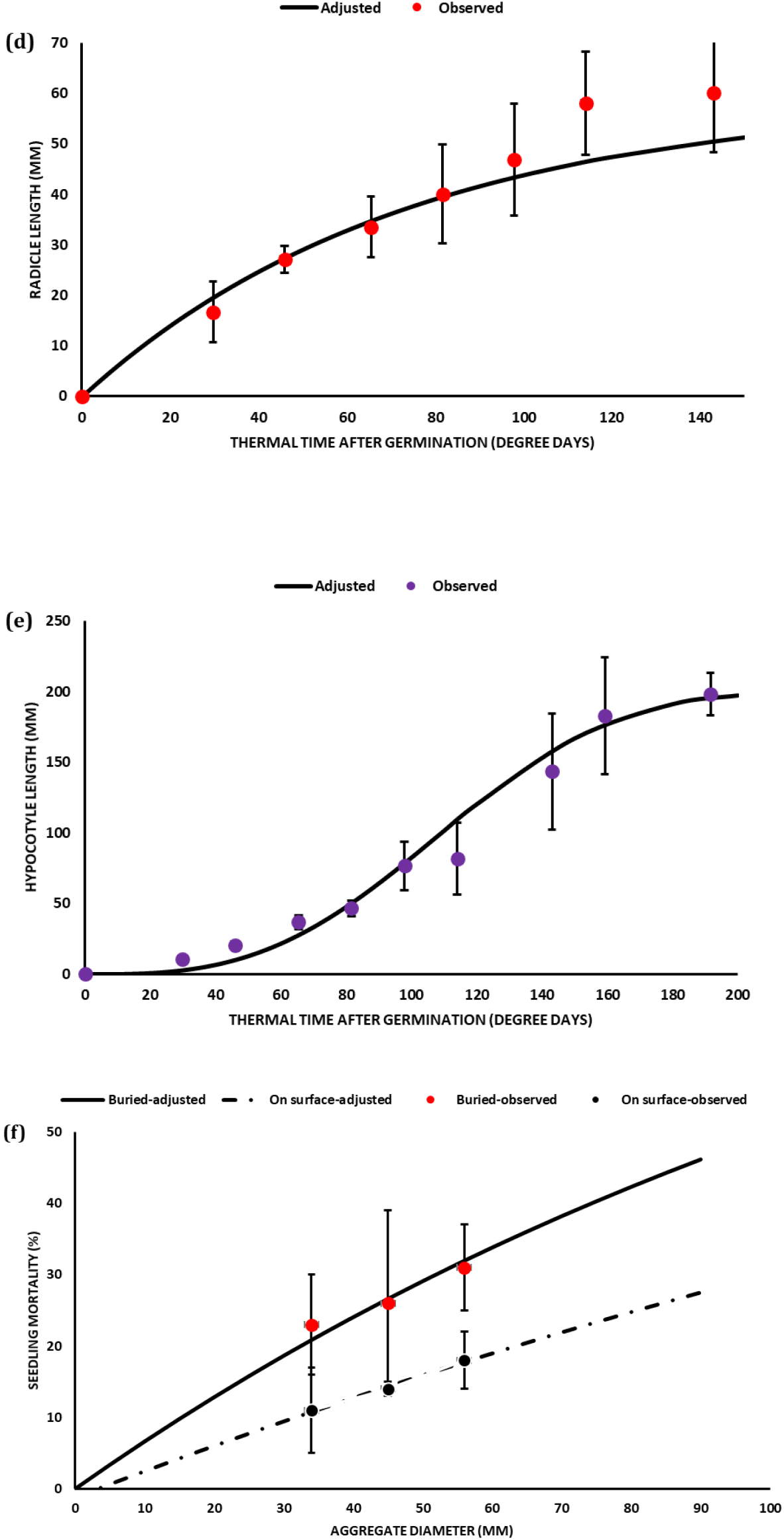
Measurement of soybean seed germination speed at 3.5, 6.5, 10, 15, 20, 25, 30, 34.5, 37 and 40 °C (a); combined values of seed germination dynamics obtained in the linear part of Figure 1a (i.e. observed at 10, 15, 20 and 25 °C) in relation to degree days (b); seed water potential (c); radicle (d), and hypocotyl (e) elongation; and seedling mortality under different soil aggregate sizes and spatial distribution (f) at 20 °C under laboratory conditions. Vertical bars reported in the figure represent standard deviations while 1/T20, 1/T40, 1/T50, 1/T60, 1/T80, and 1/T90 indicate germination speed to reach 20, 40, 50, 60, 80, and 90% germination, respectively.

When PEG 8000 was used, the rate of seed germination was 98%, 94% and 48% at water potential of −0.10, −0.25 and −0.50 MPa, respectively. No seeds germinated at water potential of −0.75, −1 and −1.50 MPa. The germination speed decreased with increasing water potential. The base water potential of the genotype ranged from −0.50 to −0.75 with an average value corresponding to −0.67 MPa **(Fig. 1c)**.

#### 3.1.2. Radicle and hypocotyl elongation under laboratory conditions

The maximum length of the radicle in heterotrophic conditions was 60 mm. Following 50% germination, 143 °Cd required for radicles to reach the maximum length **(Fig. 1d).** The radicle elongation rate during the two days following germination was 0.58 mm °Cd^−1^. The hypocotyl required 67 °Cd to reach a length corresponding to a common sowing depth (i.e. 30 mm). Results were fitted with good efficiency to a Weibull function **(Fig. 1e, Table 1)**.

#### 3.1.3. Seedling mortality under clods

When aggregates of 34, 45 and 56 mm were completely buried, 23, 26 and 31% of seedlings were blocked, respectively. When the aggregates of the same size were laid on the soil surface, 11, 14 and 18% of seedlings were blocked (**Fig. 1f; Table 1**). The rate of non-emergence due to seedlings blocked under clods was higher when aggregates were buried in the soil compared with those laid on the soil surface, especially for aggregates having higher diameter.

### 3.2. Field experiments

#### 3.2.1. Seedbed characterization

The obtained seedbed was quite coarse. In terms of number, no soil aggregate >30 mm was found either on the seedbed surface or under partially buried condition. There were 0 and 2 aggregates of 20-30 mm, and 4 and 13 aggregates of 10-20 mm on seedbed surface and under partially buried conditions, respectively. We found 1, 5, 9, 33 and 217 soil aggregates >50, 40-50, 30-40, 20-30 and 10-20 mm, respectively under completely buried conditions.

In terms of percentage of aggregate mass of the seedbed, 2, 9, 9 and 13% of the soil aggregates had >50, 40-50, 30-40, 20-30 mm. This means that over 33 % of the soil aggregate masses in the seedbed had >20 mm.

Sowing depth was highly variable which ranged from 2 to 4.5 cm. Over 60% seeds were located at sowing depth between 2.5 and 3.2 cm while other seeds were found at lower or higher soil horizons.

#### 3.2.2. Weather data

The sowing conditions were hot and dry. Daily mean seedbed climate data from sowing to emergence are reported in **Figure 2a**. The seedbed temperature ranged from 18 to 25°C at 3 cm depth and 19 to 25°C at 5 cm depth. The mean seedbed humidity ranged from 9 to 18% at 3 cm depth and 16 to 19% at 5 cm depth. The seedbed humidity at the upper soil layer showed a high variability but this variability markedly reduced with increased soil depth. At sowing depth, soil humidity was low at sowing and until the first rainfall. Most of the rainfall occurred on 5, 6, 13 and 14 das. The measured and fitted data showing the relationship between the seedbed water potential and the water content are reported in **Figure 2b**. The threshold at which the water stress began in the seedbed corresponded to 12%.

**Figure 2.**
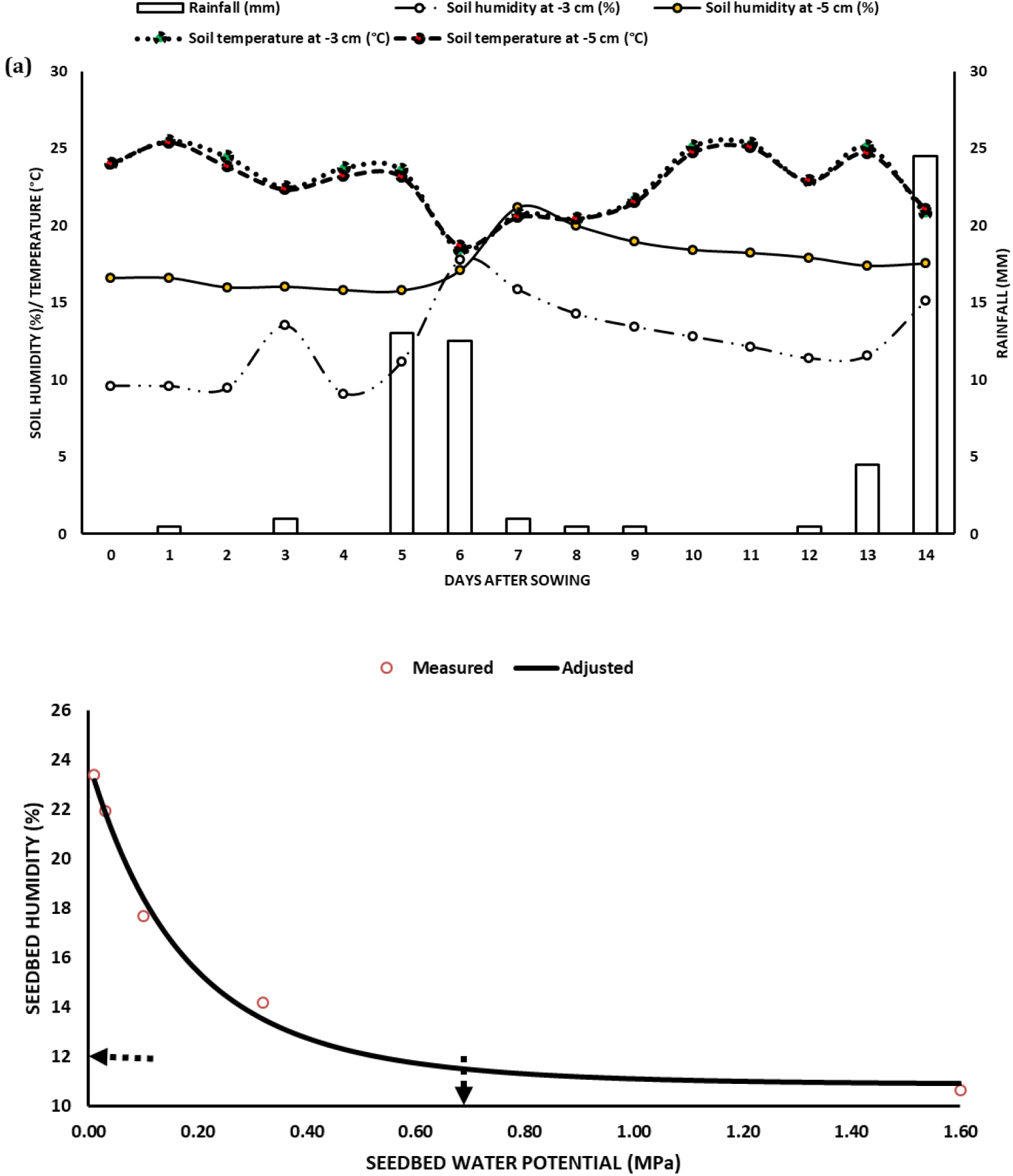
Dynamics of the seedbed soil temperature and the seedbed soil humidity and rainfall at the study site until two weeks after sowing (a), and measured and fitted soil water characteristic curves -- showing relationship between the seedbed water potential and the water content (b) -- based on the model of van Genuchten (1980). The vertical arrow on the x-axis represents the seed water potential while the horizontal arrow on the y-axis indicates threshold at which the water stress began in the seedbed that corresponded to 12%.

#### 3.2.3. Seed germination and seedling emergence under field conditions

Seed germination and seedling emergence dynamics under field condition are reported in **Figure 3.** Germination was slow under field conditions, which took 8 das to reach the maximum rate (i.e. 100%). Seedling emergence started at 5 das (i.e. 111 °Cd) and reached the maximum level (i.e. 88%) at 14 das (i.e. 261 °Cd) showing a gradual increase in emergence rate over time.

**Figure 3.**
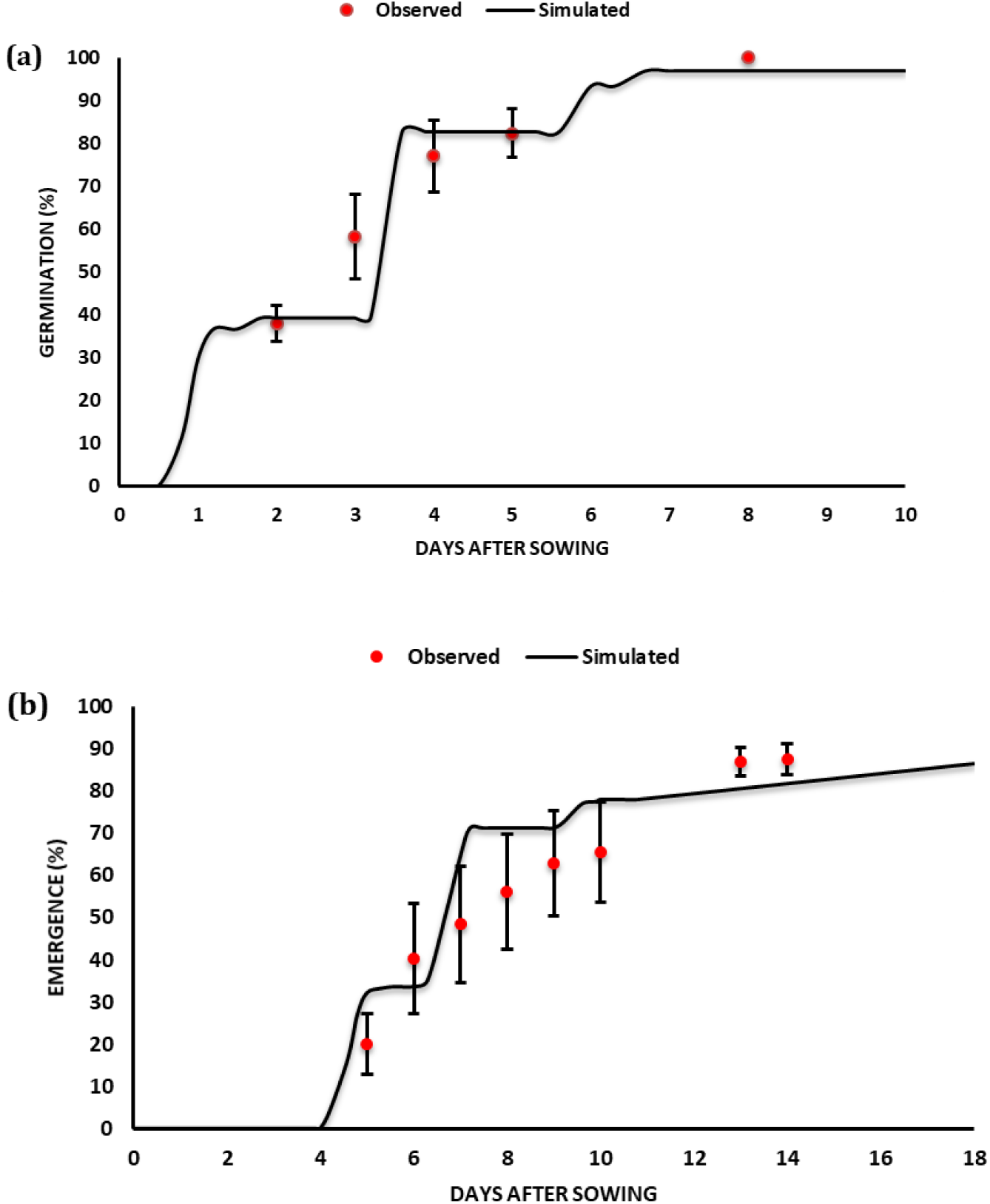
Simulated and observed values of soybean seed germination and seedling emergence rates at Auzeville experimental site, Toulouse in 2018. Vertical bars reported in the figure represent standard deviations.

#### 3.2.4. Causes of non-emergence

All the observed seeds were germinated. There were 12% losses in seedling emergence. Almost 11 % of seedling death was due either to soil clods or to a soil surface crust that was difficult to distinguish due to heavy rainfall events after sowing that created seedbed soil compaction. Soil-borne pests and pathogens caused 0.55 and 0.40% seedling emergence failure, respectively.

### 3.3. Model evaluation

Predicted vs. observed germination and emergence rates are reported in **Figure 3.** The model finely predicted germination rate, over germination courses compared with the observed dataset. The predicted final germination rate matched with the observed dataset (98.6 predicted vs. 100% observed). For germination, the model showed a relatively good fitting quality with EF of 0.65, RMSE of 0.10, and CRM of 0.04. The model prediction was very good also for emergence rate with the final predicted emergence rate almost exactly the same to the observed values (i.e. 87 predicted vs. 88% observed). For emergence, the model showed even a better fitting quality obtaining EF of 0.70, RMSE of 0.11, and CRM of −0.08.

The time to reach the maximum germination rate was the same for both predicted and observed values (i.e. 8 das) and the predicted time to reach the maximum emergence rate was also quite close (i.e. 18 das predicted vs. 14 das observed). The model also well-predicted causes of non-emergence, which was due to non-germinated seeds (1.4% predicted vs. 0% observed) and non-emerged seedlings (13 % predicted vs. 12% observed). Predicted causes of non-emergence were seedling mortality under clod (13%), that was consistent with field observations although it was not possible in the field to distinguish seedlings blocked under clods or crust (a total of 11% emergence losses) as all the seedbed was compacted -- followed by non-germinated seeds (1.4%).

### 3.4. Simulation study over 2020-2100

#### 3.4.1. Emergence rate, duration and frequency

Results on emergence rate, duration and frequency are reported in **Table 2.** Mean emergence rates ranged from 72-78% for mid March sowing, from 66-76% for 1^st^ April sowing, and from 61-76% for mid-April sowing depending on the 20-year period. No differences statistically significant were observed in terms of average emergence rate among periods and sowing dates. The frequency of poor emergence rate (<50%) ranged from 0 to 25% depending on sowing date × period and this frequency increased with later sowing dates and especially over periods. We did not find any significant interaction effect of sowing date x period on emergence rate.

**Table 2.** Emergence rate and duration (means ± standard deviation) and frequencies with <50% emergence rate and >28 days to reach the maximum emergence when analyzed by sowing date for each 20-year period and their interaction

| Sowing date | Period | Emergence (%) | Frequency (%) of emergence rate <50% | NEmax (days) | Frequency (%) of NEmax >28 days |
| --- | --- | --- | --- | --- | --- |
| Mid-March | 2020-2040 | 78<sup>a</sup> ± 13 | 0 | 37<sup>b</sup> ± 10 | 86 |
| | 2041-2060 | 73<sup>a</sup> ± 17 | 10 | 34<sup>ab</sup> ± 13 | 70 |
| | 2061-2080 | 76<sup>a</sup> ± 19 | 5 | 28<sup>a</sup> ± 7 | 65 |
| | 2081-2100 | 72<sup>a</sup> ± 15 | 10 | 28<sup>a</sup> ± 9 | 55 |
| 1<sup>st</sup> April | 2020-2040 | 71<sup>a</sup> ± 21 | 10 | 26<sup>a</sup> ± 7 | 43 |
| | 2041-2060 | 76<sup>a</sup> ± 15 | 10 | 26<sup>a</sup> ± 7 | 30 |
| | 2061-2080 | 66<sup>a</sup> ± 27 | 20 | 24<sup>a</sup> ± 8 | 25 |
| | 2081-2100 | 75<sup>a</sup> ± 20 | 5 | 21<sup>a</sup> ± 8 | 15 |
| Mid-April | 2020-2040 | 76<sup>a</sup> ± 17 | 10 | 20<sup>a</sup> ± 5 | 5 |
| | 2041-2060 | 75<sup>a</sup> ± 20 | 10 | 22<sup>a</sup> ± 7 | 20 |
| | 2061-2080 | 61<sup>a</sup> ± 27 | 25 | 21<sup>a</sup> ± 7 | 15 |
| | 2081-2100 | 70<sup>a</sup> ± 19 | 20 | 18<sup>a</sup> ± 7 | 15 |
| **Sowing dates**<br>**x Periods** | **Df** | 6 |  | 6 |  |
|  | **Significance level** | NS |  | NS |  |

Means followed by the same letter are not significantly different at $P < 0.05$ ; NS: not significant

Mean NEmax ranged from 28-37 das for mid-March sowing, from 21-26 das for 1^st^ April sowing, and from 18-22 das for mid-April sowing depending on the 20-year period. Mean NEmax values not only decreased with later sowing dates but also over periods, by more than one week for the earlier sowing dates and to a lower extent for later sowings. Mean NEmax values did not significantly decrease among periods, except for the mid-March sowing. The frequency of high NEmax (>28 days) ranged from 5 to 86% among sowing dates and periods, and it was higher for earlier sowing dates and for earlier periods.

#### 3.4.2. Causes of non-emergence rates and frequencies

Simulation results describing the main causes of non-emergence and their frequencies are reported in **Table 3**. Seedling death due to clod, seedling death under soil surface crusting, non-germination and seedling death due to drought were main causes in decreasing order of importance. Simulation outcome on the effect of sowing dates for each period and their interactions on germination rate, duration and frequency are reported in **Table 4.**

**Table 3.** Rates (means ± standard deviation) and frequencies (%) of non-emergence causes of seedlings as analyzed by sowing date for each 20-year period and their interaction. Only causes with a high frequency that could pose risks of crop emergence failure were considered which included frequency of non-germination >25% and frequency of seedling mortality due to clod, crust and drought, each >15%.

| Sowing date | Period | Non-Germination (%) | Frequency (%) of NG>25% | Average rate of SMC (%) | Frequency (%) of SMC >15% | Average rate of SMSC (%) | Frequency (%) of SMSC >15% | Average rate of SMD (%) | Frequency (%) of SMD >15% |
| --- | --- | --- | --- | --- | --- | --- | --- | --- | --- |
| Mid-March | 2020-2040 | 2<sup>a</sup> ± 0 | 0 | 13<sup>a</sup> ± 1 | 0 | 6<sup>a</sup> ± 13 | 19 | 0<sup>a</sup> ± 0 | 0 |
| | 2041-2060 | 2<sup>a</sup> ± 1 | 0 | 13<sup>a</sup> ± 1 | 0 | 10<sup>a</sup> ± 15 | 30 | 2<sup>a</sup> ± 9 | 5 |
| | 2061-2080 | 7<sup>a</sup> ± 21 | 5 | 13<sup>a</sup> ± 3 | 10 | 4<sup>a</sup> ± 9 | 10 | 1<sup>a</sup> ± 3 | 0 |
| | 2081-2100 | 2<sup>a</sup> ± 1 | 0 | 14<sup>a</sup> ± 1 | 0 | 11<sup>a</sup> ± 15 | 30 | 2<sup>a</sup> ± 5 | 10 |
| 1<sup>st</sup> April | 2020-2040 | 6<sup>a</sup> ± 20 | 5 | 13<sup>a</sup> ± 3 | 5 | 10<sup>a</sup> ± 14 | 29 | 0<sup>a</sup> ± 1 | 0 |
| | 2041-2060 | 2<sup>a</sup> ± 0 | 0 | 14<sup>a</sup> ± 1 | 0 | 5<sup>a</sup> ± 11 | 15 | 3<sup>a</sup> ± 10 | 5 |
| | 2061-2080 | 11<sup>a</sup> ± 29 | 10 | 12<sup>a</sup> ± 4 | 5 | 7<sup>a</sup> ± 13 | 20 | 3<sup>a</sup> ± 9 | 5 |
| | 2081-2100 | 7<sup>a</sup> ± 20 | 5 | 13<sup>a</sup> ± 3 | 0 | 5<sup>a</sup> ± 10 | 15 | 0<sup>a</sup> ± 1 | 0 |
| Mid-April | 2020-2040 | 2<sup>a</sup> ± 0 | 0 | 13<sup>a</sup> ± 1 | 5 | 6<sup>a</sup> ± 13 | 19 | 2<sup>a</sup> ± 7 | 10 |
| | 2041-2060 | 2<sup>a</sup> ± 0 | 0 | 14<sup>a</sup> ± 1 | 10 | 3<sup>a</sup> ± 10 | 10 | 6<sup>a</sup> ± 18 | 10 |
| | 2061-2080 | 12<sup>a</sup> ± 30 | 10 | 12<sup>a</sup> ± 4 | 10 | 12<sup>a</sup> ± 16 | 35 | 4<sup>a</sup> ± 12 | 5 |
| | 2081-2100 | 3<sup>a</sup> ± 5 | 0 | 14<sup>a</sup> ± 1 | 15 | 9<sup>a</sup> ± 15 | 30 | 4<sup>a</sup> ± 9 | 10 |
| Sowing dates<br>X Periods | Df | 12 |  | 12 |  | 12 |  | 12 |  |
|  | Significance level | NS |  | NS |  | NS |  | NS |  |

Means followed by the same letter are not significantly different at $P < 0.05$ ; NS: not significant; NG: non-germination; SMC: seedling mortality due to clod; SMSC: seedling mortality due to soil crust; SMD: seedling mortality due to drought

**Table 4.** Germination rate and duration (means ± standard deviation) and frequencies with <75% germination rate and >14 days to reach the maximum germination when analyzed by sowing date for each 20-year period and their interaction

| Sowing date | Period | Germination (%) | Frequency (%) of germination rate <75% | NGmax (days) | Frequency (%) of NGmax >14days |
| --- | --- | --- | --- | --- | --- |
| Mid-March | 2020-2040 | 98<sup>a</sup> ± 0 | 0 | 16<sup>a</sup> ± 7 | 67 |
| | 2041-2060 | 98<sup>a</sup> ± 1 | 0 | 15<sup>a</sup> ± 8 | 40 |
| | 2061-2080 | 93<sup>a</sup> ± 21 | 5 | 15<sup>a</sup> ± 6 | 35 |
| | 2081-2100 | 98<sup>a</sup> ± 1 | 0 | 13<sup>a</sup> ± 7 | 40 |
| 1<sup>st</sup> April | 2020-2040 | 94<sup>a</sup> ± 19 | 5 | 11<sup>a</sup> ± 5 | 24 |
| | 2041-2060 | 98<sup>a</sup> ± 0 | 0 | 11<sup>a</sup> ± 5 | 25 |
| | 2061-2080 | 89<sup>a</sup> ± 29 | 10 | 12<sup>a</sup> ± 8 | 35 |
| | 2081-2100 | 93<sup>a</sup> ± 20 | 5 | 11<sup>a</sup> ± 6 | 30 |
| Mid-April | 2020-2040 | 98<sup>a</sup> ± 0 | 0 | 9<sup>a</sup> ± 6 | 14 |
| | 2041-2060 | 98<sup>a</sup> ± 0 | 0 | 11<sup>a</sup> ± 8 | 30 |
| | 2061-2080 | 88<sup>a</sup> ± 30 | 10 | 11<sup>a</sup> ± 7 | 25 |
| | 2081-2100 | 97<sup>a</sup> ± 5 | 0 | 8<sup>a</sup> ± 8 | 20 |
| **Sowing dates x Periods** | **Df** | 6 |  | 6 |  |
|  | **Significance level** | NS |  | NS |  |

Means followed by the same letter are not significantly different at $P < 0.05$ ; NS: not significant

##### 3.4.2.1. Non-germination

The variability in mean non-germination rate ranged from 2 to 12% depending on the period and sowing date. The sowing date x period interaction effect on non-germination rate was not statistically significant (p<0.05). The frequency of high non-germination (>25%) ranged from 0 to 10% **(Table 3)**. The main causes for non-germination were low temperatures for early sowings and drought for later sowings.

Average NGmax values ranged from 8 to 16 days, which generally tended to decrease with later sowing dates and periods **(Table 4)**. The frequency of high NGmax (>14 days) ranged from 14 to 67%, which also generally decreased with later sowings and over the periods. There was no statistically significant effect (p<0.05) of sowing date, period and their interaction on mean NGmax values.

##### 3.4.2.2. Seedling mortality due to clods

There was no variability in seedling mortality rate due to clods among the sowing dates or periods, which ranged from 12 to 14% **(Table 3)**. This little variability was due to the same seedbed structure used for all simulations, without any influence of the sowing dates and periods. No statistically significant effect of the sowing date x period interaction was observed on seedling mortality rate due to clods.

##### 3.4.2.3. Seedling mortality due to crust

The variability of seedling mortality rate due to soil surface crust ranged from 3 to 12% **(Table 3).** Because soil surface crust prevents emergence only when it becomes dry with soil evaporation, this effect was observable for mid-April sowing, particularly for the last two periods characterized by much higher seedling mortality rate compared with the first two periods. There was no significant effect (p<0.05) of sowing date, period or their interaction on seedling mortality rate due to crust. The frequency of high seedling mortality rate due to soil surface crust (>15%) ranged from 10 to 35% and was higher for the 2081-2100 period for all but 1^st^ April sowing **(Table 3)**.

##### 3.4.2.4. Seedling mortality due to drought

Average seedling mortality rate due to drought ranged from 0 to 6% **(Table 3)**. No significant effect of sowing date, period or their interaction was found on seedling mortality rate due to drought. The frequency of high mortality due to drought (>15%) ranged from 0 to 10% and was higher for the latest sowing date and it also increased with periods. Some seedling mortality due to drought was observed even for the earliest sowings.

## 4. Discussion

### 4.1. Observed and predicted germination and emergence dynamics

This study has generated reference values related to soybean seed germination and seedling emergence. This makes possible a comparison of soybean with a set of other leguminous crops and cover crops, for which the same variables were measured and values have been reported in the literature (Dürr et al., 2015; Tribouillois et al., 2016). We did determine the base temperature for seed germination but not for hypocotyl elongation because as the temperature response for germination and development of other plant organs has been reported to be the same for several crop species (Parent and Tardieu, 2012). The comparison among species shows that leguminous crops have a wide range of temperature and water potential threshold values for germination, and this information can help while making decisions related to cropping system adaptation to changing climate or new environments. We found that the base water potential value of soybean is high (−0.67 MPa), close to that of barrel clover (−0.7 MPa), and other cover crop species including grass pea (−0.8 MPa), wild lentil (−0.7 MPa), hop (−0.6 MPa), yellow sweet (−0.7 MPa), Egyptian clover (−0.8 MPa) and crimson (−0.8 MPa) clovers, common sainfoin (−0.6 MPa), fenugreek (−0.5 MPa) and purple vetch (−0.7 MPa). However, the base water potential value of soybean is higher than other leguminous crops such as pea (−2.2 MPa), common bean (−2.3 MPa), lupin (−1.5 MPa) and chickpea (−1.8 MPa), and cover crops including common (−1.1 MPa), and winter (−1.1 MPa) vetches, highlighting that soybean is very sensitive to water stress in the seedbed compared to other legume crops.

The base temperature for germination of soybean was 4°C, an intermediate value close to that of some cover crop species such as grass pea (3.5°C), fenugreek (4.2 °C), and common vetch (4.1 °C). Nevertheless, this base temperature for germination of soybean was lower than that of common bean (8°C), cowpea (8.5 °C), mungbean (10°C), and other cover crop species including Egyptian (6.1 °C) and crimson (6.4 °C) clovers; while it was higher than that of faba bean (0.4°C), chickpea (0°C), lentil (2°C), lupin (−0.8°C) and pea (−1°C), and other cover crop species including wild lentil (0.8 °C), blue lupin (0.8 °C), hop clover (0.6 °C) and yellow sweet (0.8 °C) clovers, common sainfoin (0 °C), purple (2.1 °C) and winter (1.4 °C) vetches.

Time to reach mid-germination was 17°Cd for soybean suggesting how fast this species germinates when water is not a limiting factor. The time to mid-germination of soybean was lower than many other leguminous crops including pea (24-34 °Cd), lentil (21-25 °Cd), chickpea (45°Cd), cowpea (27°Cd), and fababean (47°Cd), while it was much closer to that of common bean (14°Cd) and mungbean (12°Cd).

The optimum temperature for soybean germination was 30°C, which is higher than that of pea (22°C), lentil (24°C), and faba bean (25°C) and many other cover crop species such as grass pea (27°C), blue lupin (26°C), hop (26°C), yellow sweet (25°C), and crimson (26°C) clovers, common sainfoin (24°C), purple (24°C), common (22°C) and winter (20°C) vetches. The optimum temperature for soybean germination is close to that of fenugreek (30°C), Egyptian clover (30°C) and also wild lentil (32°C) and chickpea (32.5°C), but lower than that of common bean (32-34°C), cowpea (35°C), and mungbean (40°C).

While soybean germination rapidly occurred under laboratory conditions (ca. 2 days or 35°Cd) with no temperature or water stress, time to reach the maximum germination was much longer (8 days or 169°Cd) in our field experiment. Despite an optimal seedbed temperature, the slow germination was due to the low water content in the seedbed that prevented seeds from being readily germinated. Only the first rainfall at 5 das allowed to complete germination of seeds. Also the emergence was very slow in the field, despite an optimum seedbed temperature, due always to water stress in the seedbed. Comparison of the predicted germination and emergence courses and final rates showed only little differences between observed and predicted values. This clearly highlights the robustness of the prediction quality of the model which was also reported previously (Constantin et al., 2015; Dürr et al., 2016) and the good estimates of the parameters values obtained for germination and growth via the laboratory experiments.

### 4.2. Simulation study under future climate change

A previous study (Lamichhane et al 2019) showed that, for early spring sowings, climate change, simulated under the IPPCC 8.5 scenario, ie the most pessimistic, will become more significant after 2060 in Northern France, with progressively increasing mean seedbed temperatures by +2 °C in February, March and April after 2060. There will be a higher variability of rainfall as well, although with no overall change of its cumulated values with the exception for the mid-April sowing. Indeed, for this latter sowing date, the simulated average cumulated rainfall will be almost two-fold lower compared with the 2000–2018 period. These climatic data were used to feed the SIMPLE model and run the present simulation study on soybean establishment. This study showed no strong risks for a successful establishment of soybean in Northern France in the coming decades for the simulated sowing dates under the chosen scenario. A previous study (Lamichhane et al, 2019), that considered the same periods took into account in this study, highlighted that the 2020 – 2040 period (will be very close to the current climate and that important changes in climatic conditions will occur only after 2040. The average simulated soybean emergence rates for the period 2020-2040 were 71-78% depending on the sowing date. Soybean has a rather low base temperature for germination and it can readily germinate and emerge, as long as there is water availability in the seedbed. However, emergence time is much delayed for the earlier sowings that may provide opportunities for pathogens to attack germinating seeds and emerging seedlings.

In terms of the percentage of seedling losses, seedling mortality rates under clods or soil surface crusting were the most frequent. Germination stage was less impacted than emergence by the abiotic factors, although too low temperature with earlier sowings or drought during the later sowings slightly affected the seed germination process. Taking together both the germination and seedling losses, water stress however can have a quite large effect on emergence results, especially for the mid-April simulated sowing.

Seed germination and seedling emergence rates of soybean simulated by the SIMPLE crop emergence model could be overestimated because this model does not take into account the effect of soil-borne pests and pathogens. Nevertheless, stand losses due to these biotic stresses could be still limited under current cropping practices. While damages due to *Rhizoctonia solani*, one of the most important soil-borne pathogen causing damping-off, have been sporadically reported on soybean (TerresInovia, 2018), biotic stresses are not still a constraint for soybean crop establishment in France. Indeed, currently, soybean is grown in France without chemical seed dressing also because this practice is often not compatible with soil or seed inoculation of bacteria that promote nodulation (Campo et al., 2009; Zilli et al., 2009). Another reason explaining the absence of biotic stresses on soybean is that this crop is still grown on a small surface in France and it is often introduced in diversified cropping systems -- especially rotations with maize and wheat— once every five to six years (Lecomte and Wagner, 2017). Moreover, the most important soybean basin is Southwestern France where cool and moist seedbed conditions, favorable for soil-borne pathogens, rarely occur. Indeed, field diagnosis on the causes of non-emergence further confirmed that soil-borne pests and pathogens rarely affect soybean stand. However, the situation may change in the future as farmers may tend to perform early sowings to escape from summer drought and save irrigation water (Maury et al., 2015). Another reason for early sowings could be that this practice allows growing late season varieties that are generally more productive than the early season ones. On the other hand, introducing soybean more frequently in the rotation could increase risks of soil-borne pathogens, especially without chemical seed dressing, although this is yet to be investigated in future studies. In the last five years, there is an exponential increase in surface grown to soybean (approximately 15000 ha/year; FAOSTAT 2018) and this crop could expand to the Northern regions of France, characterized by higher risks of biotic stresses. Stand losses due to biotic stresses have been reported from Northern Europe such as Belgium (Pannecoucque et al., 2018). Likewise, soybean is one of the crops subjected to heavy attacks from soil-borne pathogens during its crop establishment phase across countries such as USA (Ajayi-Oyetunde and Bradley, 2016; Rojas et al., 2016), the major producer of soybean worldwide. In the US, frequent returns of this crop into the same field and rotations with crops (especially maize) subjected to attacks from the same soil-borne pathogens have been reported as key drivers of this disease pressure.

Seedling predation due to birds, especially for early sowings can also be a serious problem and seedling losses can reach 100% if no protection measures are applied during the emergence phase.

## 5. Conclusions

Soybean is an interesting leguminous crop at the global level both from socio-economic and environmental points of view. Nevertheless, this crop still does not occupy the place it would merit in the European cropping systems. Relevant information on this crop including key characteristics of the seeds and seedlings, as well as biotic and abiotic factors affecting its establishment are still poorly known, hindering exploration of suitable areas for soybean cultivation. Therefore, more research on soybean in general, and on factors affecting its stand development in particular, can contribute to increased soybean cultivation in the EU. To this objective, this study is a first attempt to determine the quality of soybean establishment not only in a region with previous cropping history, but also to explore new areas that were not a historical growing basin of this crop, but would become suitable for its establishment under future climate change.

Results of the field observations in this study showed that soybean sowing under sub-optimal conditions (coarse seedbed, late sowing date, no irrigation before the crop establishment etc.) resulted favorable in Southwestern France in terms of crop stand establishment. Results of field observation were confirmed by simulation that took into account the most pessimistic future climate change scenario in northern European climatic region. Therefore, risks of poor stand establishment seem to be relatively low for soybean due to the capacity of this crop to promptly germinate when water is not a limiting factor. While the RCP weather scenario considered in this study was the most pessimistic from climate change point of view, it actually did not result that negative for soybean establishment. In contrast, the less pessimistic RCP scenarios could result less favorable for soybean establishment given that lower average temperature of the seedbed will mean higher number of days needed for soybean germination and emergence, with higher risks of attacks from biotic factors dwelling in the soil. Given that soybean is particularly sensitive to water stress in the seedbed, growers are encouraged to perform sowings at the beginning of April, i e earlier than the currently practiced sowing date. Nevertheless, field access could be a limiting factor to perform early sowings, especially across northern European regions, due to higher water content of the soil top layers (Lamichhane et al 2019). Therefore, farmers should find a trade-off and perform early sowing as soon as the soil humidity allows them to enter into the field with agricultural equipments.

Such results was obtained by using local daily predicted data for future climate and crop-model simulations. Despite all the limits of these models, such studies contributes to better understand and anticipate the future possible changes in cropping systems. Overall, our results are encouraging for farmers who will be willing to perform early sowing in order to adapt to ongoing climate change. Farmers in France and Northern Europe, who currently do not grow soybean or hesitate perform sowing are encouraged to consider soybean to diversify their cropping systems.

## Acknowledgements

The authors thank the technicians of the Vasco research team, UMR AGIR, and those of the Auzeville experimental unit for field crops, for their kind support during this study. We are grateful to Gilles Tison (UE Auzeville, INRA) for his availability to discuss and program field experiment and Christine Le Bas (INRA, Orléans) for her suggestions concerning methods to determine the soil water potential. This study was funded by a starter grant of the INRA’s Environment and Agronomy Division to the first author.

